# A model for the spatio-temporal design of gene regulatory circuits

**DOI:** 10.1101/522946

**Authors:** Ruud Stoof, Alexander Wood, Ángel Goñi-Moreno

## Abstract

The design of increasingly complex gene regulatory networks relies upon mathematical modelling to link the gap that goes from conceptualisation to implementation. An overarching challenge is to update modelling abstractions and assumptions as new mechanistic information arises. Although models of bacterial gene regulation are often based on the assumption that the role played by intracellular physical distances between genetic elements is negligible, it has been shown that bacteria are highly ordered organisms, compartmentalizing their vital functions in both time and space. Here, we analysed the dynamical properties of regulatory interactions by explicitly modelling spatial constraints. Key to the model is the combined search by a regulator for its target promoter via 1D sliding along the chromosome and 3D diffusion through the cytoplasm. Moreover, this search was coupled to gene expression dynamics, with special attention to transcription factor-promoter interplay. As a result, promoter activity within the model depends on its physical separation from the regulator source. Simulations showed that by modulating the distance between DNA components in the chromosome, output levels changed accordingly. Finally, previous experimental results with engineered bacteria in which this distance was minimized or enlarged were successfully reproduced by the model. This suggests that the spatial specification of the circuit alone can be exploited as a design parameter to select programmable output levels.

## 1 Introduction

The rational engineering of biological systems involves the design and implementation of gene regulatory circuits to turn input signals into output responses (*1*, *2*). Inspired by electrical engineering many such circuits have been successfully built in bacteria, including oscillators (*3*, *4*), switches (*5*, *6*), Boolean logic gates (*7*), counters (*8*) or multiplexers (*9*). Unlike electronic circuits in which signals are carried from one component to the next trough dedicated and insulated wires, genetic circuits share the same physical space for all communications. This is to say, molecular signals share the same *wire* (i.e. the entire volume of the cell) which is complex, crowded and dynamic (*10*, *11*). The inability to separate the genetic circuits from the rest of the intracellular milieu makes gene regulation an intrinsically space-dependent process. Each transcription factor (TF) that interacts with a promoter to regulate gene expression must *travel* some distance through that milieu from where they are expressed to their target. Despite the small volume of a bacterial cell, it has become increasingly clear in recent years that spatial constraints have important implications on bacterial functions (*12*). The analysis of spatial dynamics at the population level (e.g. inter-bacterial communication or pattern formation) is an active field (*13* –*16*). However, current synthetic biology has little regard to the spatiotemporal complexity of the *intracellular* environment - an issue that deserves further attention in order to integrate spatial constraints within the design-build-test lifecycle.

Structural analyses on bacterial cells have helped to clarify this internal complexity, where a non-compartmentalized, but highly organized chromosome is compacted (*17*). Rather than being randomly dispersed throughout the cell, the chromosome is organized into four large macrodomains and two *non-structured* domains (*18*, *19*). In addition to this, it is further organized into smaller, more dynamic microdomains (*20*, *21*). Such chromosomal structure is heavily linked to genetic function; it has been observed that genes which are co-regulated are often clustered and retained in close proximity, not only in terms of base pairs (*22*) but also considering the 3D folding of the chromosome (*23*, *24*). As a result of this organisation, molecular species are also somewhat localised. For instance, some TFs are not homogeneously distributed within the volume of a cell (*25*), but localised around their interacting DNA components e.g. their coding gene and cognate promoter (*26*). The coupled nature of transcription and translation in prokariotes (*27*) is theorised to produce locally high distributions of TFs near the site of their expression (*28*), making co-localization of TFs coding gene and their target promoter a potential evolutionary solution for the tight control of protein production. The physical separation of co-regulated genes is then revealed as potential key parameter for such expression control. Is has been shown that, for instance, the further away from the TF’s coding gene that the target is located in the chromosome, the less effective the regulation is (*29*). Therefore, the strength of a regulatory interaction, either repression or induction, may change with increasing or decreasing this relative distance - an issue that deserves further attention.

Intergenenic distance is also at the origin of gene expression variability, the so-called genetic noise (*30*, *31*), which in turn leads to cell differentiation (*32*). In a previous experimental work (*33*) we correlated noise patters to both [i] distance from TF source to the target promoter, and [ii] total TF numbers in the cell. When the TF total amount was decreased to basal levels, the intergenic distance became a decisive parameter for fine-tuning target noise. Such noise was high if source and target were far apart, but was almost eliminated when both DNA elements were moved into close proximity.

Mathematical and computational modelling have become fundamental tools in synthetic biology (*34*, *35*). Models can predict molecular behaviour, inform the design process by advising on the selection of DNA parts and help to understand intricate experimental results. One of the challenges for mathematical modelling in synthetic biology is to re-define abstractions, assumptions and processes as new mechanistic knowledge of molecular systems arises. However, there is currently no modelling framework that enables the simulation of regulatory interactions in a spatial context as described above. Nevertheless, there are fundamental descriptions of the movement of TFs within the intracellular volume of a cell. Facilitated diffusion modelling has shown that TFs will quickly *find* specific locations on the DNA if these are in close proximity to their point of generation (*28*, *36*–*38*). Indeed, transcription factors are at one location at a given time, which adds complexity compared to non-spatial homogeneous models. Breaking with the traditional assumptions of homogeneous models involves explicitly modeling 1D diffusion (hopping and sliding) along the chromosome, un/re-binding of TF from/to non-specific DNA regions and 3D diffusion across the cytoplasm (*39*).

Here, we develop a modelling framework to simulate regulatory interactions in such a spatio-temporal setup. Firstly, the model is described and compared against a (typical) homogeneous model. Secondly, the model is used to fit experimental data where intergenic distance was minimized or enlarged (*33*). Finally, distance is highlighted as a potential *design parameter*. That is, intergenic distance is used to fine-tune the predicted performance of synthetic circuits.

## 2 Results

Figure 1 shows the spatio-temporal principles that govern gene regulation within the model. Fundamental to this approach is the time it takes for a transcription factor (TF) to go from its encoding gene, where it is expressed, to its cognate promoter, where the TF binds. Therefore, these two regions are referred to as *source* and *target*, respectively. Since TFs must actively reach the target, the distance between them (i.e. base-pairs separation in the chromosome) is the key parameter that the model revolves around (Figure 1A). This implies that TFs will *not* be automatically available to bind the target after expression, which is the customary view of what we refer to as *homogeneous* models (*40*, *41*) (Figure 1B). Importantly, such models are inconsistent with having intracellular spatial dynamics. Specifically, the basic dynamics of TFs searching for the target need to be considered, as summarized in Figure 1C: sliding and hopping (combined in our model), 3D diffusion (or *global search* as it is explained next) and non specific un-/binding to DNA regions that are not the target.

**Figure 1:**
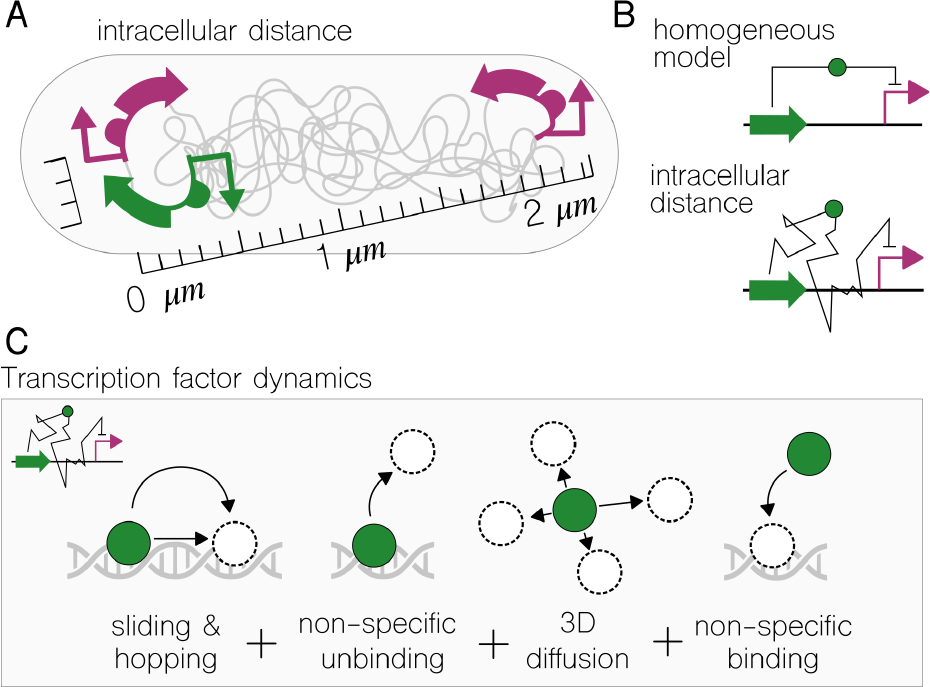
Spatiotemporal principles of gene regulation. **(A)** Diagram of three chromosomal insertions: one source of transcription factors (green) and two target regions (purple). One of the target regions is co-located with the source; the other one is spatially separated. **(B)** A Schematic comparison between *homogeneous* and *spatial* models. While the former assumes that transcription factors are *instantly* available to bind, the latter explicitly simulates the *travel* from source to target. **(C)** Transcription factor dynamics through facilitated diffusion: 1D sliding, hopping, non-specific un-/binding and 3D diffusion. The model adds these dynamics to promoter activity and gene expression events.

### Modelling the search of a transcription factor for its target

The model simulates a negative regulation where the source is formed by a constitutive promoter that expresses a downstream repressor, and the target consists of the corresponding repressible promoter upstream a reporter gene (e.g. green fluorescent protein, GFP). Spatial constraints do not affect the process of mRNA transcription, which is determined by a set of Ordinary Differential Equations (ODEs):

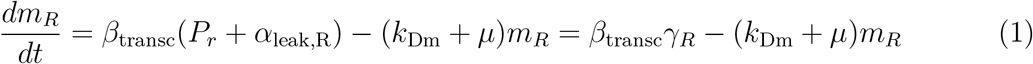

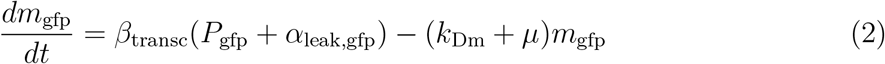

where *m*_*r*_ and *m*_gfp_ represent the amount of mRNA molecules that encodes the repressor and reporter proteins respectively, *β*_transc_ is the rate of transcription of a non-repressed target promoter, *α*_leak,i_ is the basal transcription of the target promoter when it is repressed, *k*_Dm_ the RNA degradation rate, *P*_*gfp*_ the target promoter, and *μ* the growth dilution rate. The model describes the “availibility” of the target promoter (*γ*_*R*_), with *γ*_*R*_ = 1 meaning full promoter activity and *γ*_*R*_ = 0 meaning no transcription.

Spatial effects are modelled upon mRNA translation, which implies mRNA molecules are left out of space-dependent constraints. This assumption draws on the coupling of transcription and translation in prokaryotes (*27*, *28*) along with fast mRNA decay (see Methods). Immediately after the generation of a transcription factor, this is classified as *local* and it is 1D diffusion along the neighbouring chromosome (sliding + hopping) that determines TF’s location and dynamics. After this initial period (timescale of seconds) the regulator will unbind from the chromosome into the cytosol and rapidly lose location autocorrelation due to fast 3D diffusion (see Table 1 and Methods). From that moment onward (until it degrades), the TF is classified as *global*. The ODEs that calculate the amount of both local and global TFs over time are:

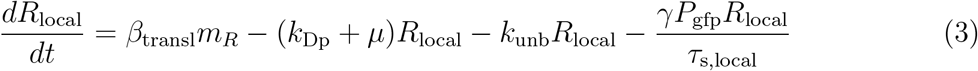

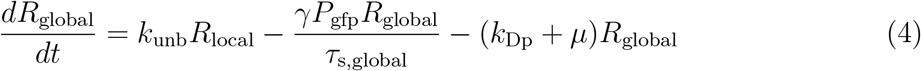

where *R*_local_ and *R*_global_ are the amount of local and global repressor, respectively, *β*_transl_ is the rate of translation from mRNA,*γ* is the fraction of TF binding when located at the TG, *k*_Dp_ is the protein degradation rate and *τ*_s,global_ the global search time. The equation for the local search time, *τ*_s,local_, is given in 10. While 1D diffusion is relatively slow compared to 3D diffusion, local search times can be orders of magnitude shorter than global ones (seconds instead of tens of minutes) if source and target are co-localised. TF location is always known during local search. However, that is not the case during global search, when TFs combines 3D diffusion and 1D sliding - their location is random in respect to the source.

**Table 1:**
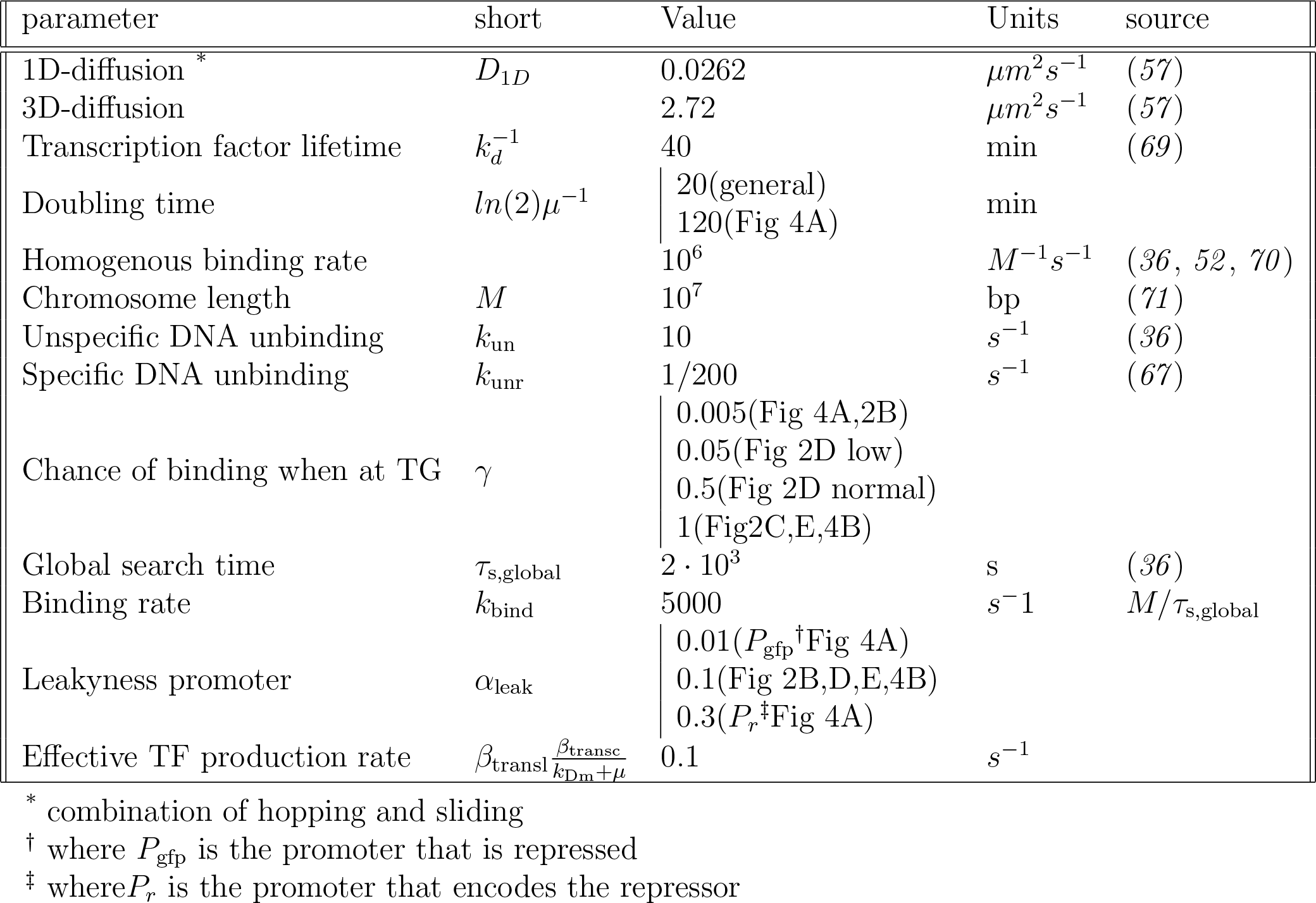
Parameters used in simulations

When, finally, a repressor binds to the target promoter, the expression of the reporter gene is inhibited. The set of ODEs that describe this is as follows:

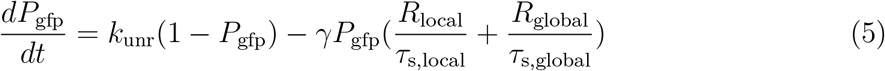

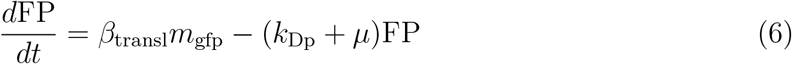

where *k*_unr_ is the specific unbinding rate of the repressor from the target promoter. The model was simplified with quasi-steady-state conditions (Supplementary File S1) and parameters were obtained from experimental values when possible (Table 1).

### System sensitivity to intergenic distance modulation

Gene expression, and more specifically promoter activity, reflects the upstream dynamics of the cognate regulatory machinery. In traditional homogeneous models, such promoter activity is determined by the overall number of TFs and a constant binding rate. In contrast to this, the addition of spatial resolution to modelling makes TF binding dependent of [i] the whereabouts of the TF (i.e. local or global), [ii] diffusion speeds, [iii] the distance between the source of TFs and the target promoter, [iv] binding rate and [v] non-specific and specific unbinding rates (Figure 2).

**Figure 2:**
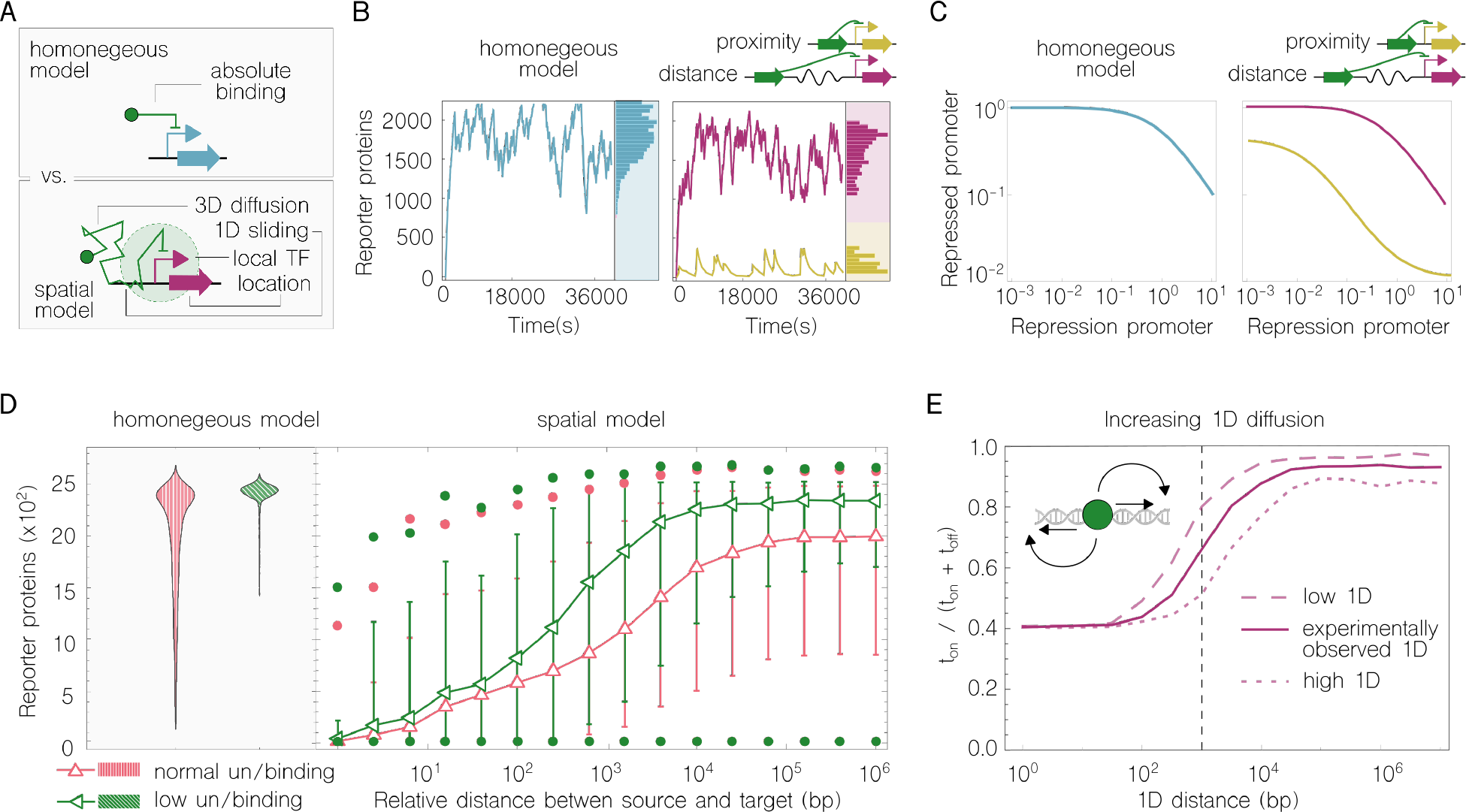
Analysis and comparison of a negative regulation (repression) in both spatial and homogeneous models. **(A)** The binding of a transcription factor to its cognate promoter is the key difference of both modelling approaches. While a single absolute value is enough in a homogeneous model, binding events depend on 1D/3D sliding/diffusion and the relative distance from TF source in spatial models. **(B)** Time course simulation of gene expression in the two models. The homogeneous simulation is similar to the distant source-target scenario (rates were pre-defined to this end in order to show the effects of decreasing distance). **(C)** Characterization of promoter activity in both source (*x* axis) and target (*y* axis) regions. Axis measure the normalized proportion of time (from 0 to 1) that a promoter is in its *active* state. If the source promoter is predominately *active* (*x* axis to the right), the repression is more efficient and the target promoter is strongly *inhibited* (*y* axis to the bottom). A sharper correlation is found on the proximity use-case. **(D)** Gene expression noise variation in relation to source-target distance. Noise was analyzed in both homogeneous and spatial models with two sets of rates for binding and unbinding. The relationship between the two simulations in the homogeneous model (left) is somewhat mimicked by the spatial model (right) when the relative distance from source to target is high. If such distance is low, the relationship is the opposite. Error bars show noise ranges within standard deviation; dots represent outlier values. **(E)** Increasing 1D diffusion decreases the impact of intergenic distance. The fraction of time the target promoter is active (i.e. without the repressor bound; *y* axis) increases as the target moves away from the source. However, increasing 1D diffusion prevents (up to a limit) this trend.

Simulations of an inhibition interaction (i.e. repression) using both homogeneous and spatial models were compared (2B). For the sake of comparison, the non-spatial parameters in both models have the same values (e.g. transcription, translation or molecule degradation). That is, source-target distance, within TF dynamics, is the only difference between the two models. Simulations returned similar expression levels for both approaches, providing that the relative location of genetic components was distant in the spatial model. However, when components were co-localized in the spatial model the time series simulations were very different. Upon co-localization, the repression was observed to be stronger (i.e. less reporter expression). A closer look at promoter availability (fraction of time it is in its active form) confirmed this trend - the activity of the source promoter has a faster (and stronger) impact on the target when placed in proximity, since the interaction is not diluted within the volume of the cell (Figure 2C). It is not coincidental that the simulations gave similar results between the homogeneous and spatial models when source-target distance was maximised. This is because space-specific parameters in the spatial model were setup to that end. Therefore, the comparison informs about what the reduction of source-target distance leads to, which is, as described above, a lower reporter expression (i.e. stronger regulation). However, at high intergenic distances, the majority of TFs reach the target promoter via global (instead of local) search, which in our model generates a uniform distribution of TFs along the chromosome. As a result, the spatial parameters that are more relevant during local searches, lose significance and the simulation converges to the homogeneous model. If space-specific parameters had been established so that the proximity scenario had matched the homogeneous model in the first place, the simulation would returned opposite results.

Gene expression noise (*42*, *43*) is decisively influenced by intergenic space separation. Nonetheless, the so-called *bursting* effect (*44*, *45*), a pulse-like expression activity that results from a transcription factor binding and unbinding its cognate promoter, is a major source of genetic noise. Given that physical source-target separation modifies the binding process through modulating the availability of TFs, it can be concluded that it has a direct influence in transcription bursts. Figure 2D shows the variation in gene expression noise in relation to source-target distance. Results are shown for two sets of binding and unbinding rates: the first one referred to as *normal* (obtained from the experimental literature) and the second one termed *low* (one tenth of the previous, Table 1). While the noise returned by the homogeneous model shows a clear trend towards a wider noise pattern in the *normal* scenario, the spatial simulation arrived at both this and the opposite trend depending on the source-target separation selected. When such separation is short, genetic noise under the *normal* (un)binding rates is narrower than using the *low* rate values. In contrast, upon increasing separation the situation reversed. This suggests that protein variability is more than the mere consequence of stochasticity and can be deterministically controlled by modulating intergenic distance alone.

### Nonlinearity of regulation response to intergenic distance

The effects of source-target separation in reporter expression do not vary in direct proportion to the increase in distance (Figure 2E). In fact, up to 10^2^ base pair (bp) separation, the effects were found to be effectively the same. A similar performance was observed when the separation was above 10^4^ bp, when further differences cannot be appreciated. However, the impact of such separation increases in all values in between, almost proportionally, with 10^3^ bp being halfway in the overall response curve. Figure 2E implies that the origin of such nonlinearity lies in the balance between local and global search: local search predominates in the first region (0-10^2^bp), within which the location of the target promoter will not make a difference; both local and global searches are combined in the second region (10^2^-10^4^bp), that shows proportional effects when increasing distance; global search predominates in the third region (10^4^-10^7^) which also resulted in a plateau-like response function. Although the thresholds between such regions can be altered by decreasing/increasing 1D diffusion (Figure 2E), the overall pattern remains the same. This suggests that spatial effects are stable at two source-target relative locations, *very close* and *very far*, while at middle points the regulation would change rapidly. The distribution of transcription factors along the chromosome is therefore central to the model.

To analyze this distribution, we examined the location of TFs during local search upon their generation from the source. Figure 3A records the trajectories of individual regulator molecules along the chromosome while performing 1D difussion (sliding + hopping) until they unbind into the cytosol. Given the coupling of transcription and translation in bacteria (*46*–*48*), it is safe to assume that newly generated regulators start their search from (nearby) the source coding region (0bp in the graph). The visualization of 1D movement gives two conclusions. First of all, the timescale for the local search is of the order of seconds - with many TFs spending less than a second bound to the local chromosome. The second conclusion is that the local region is crowded with TFs searching for their target, which means that any regulatory interaction will be stronger / more efficient within this chromosomal segment.

**Figure 3:**
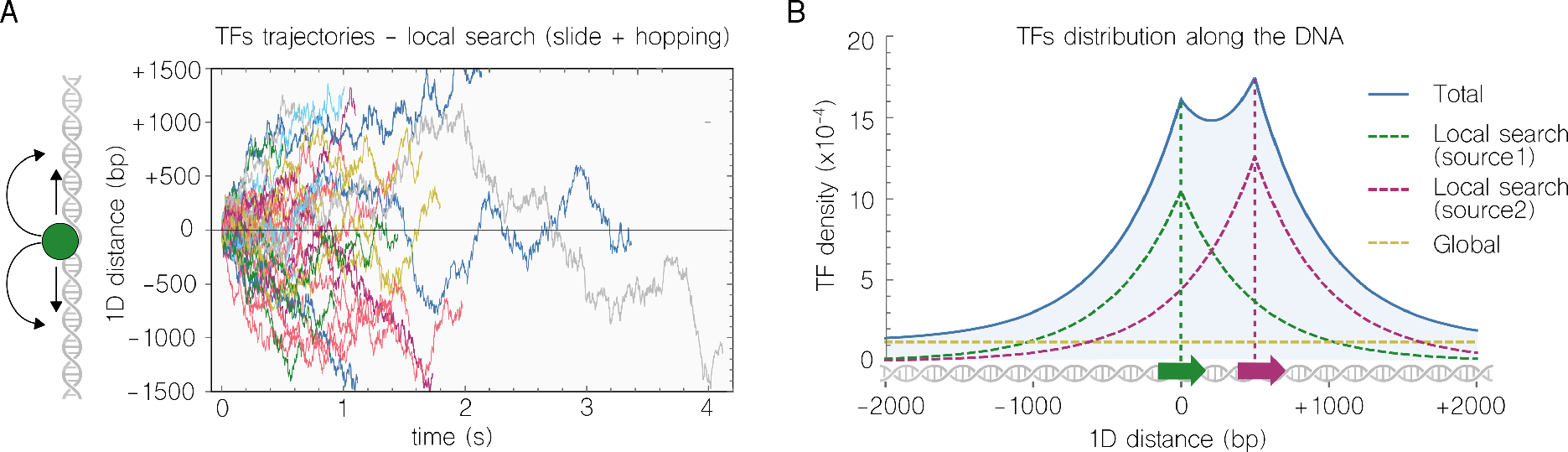
Transcription factor distribution in relation to the location of source and target regions. **(A)** Simulations of the time (*x* axis) during which individual transcription factors (coloured trajectories) slide and hop along the chromosome and around their encoding gene location (*y* axis). Each simulated line ends when the transcription factor *jumps* to 3D diffusion. **(B)** Density of transcription factors along the chromosome due to specific modelling dynamics. Global search (yellow line) results in a flat distribution where, due to fast 3D diffusion, the transcription factors are equally spread along the cell’s volume. Local search around source (green line) and target (purple line) regions favours the accumulation of transcription factors in this areas due to expression and repression, respectively. Total distribution coloured in blue.

When the total density of TFs along the chromosome were calculated (Figure 3B), it was observed that such higher local concentration of TFs could also be found in the neighbourhood of the target. While global search generated a uniform distribution (i.e. same value for all chromosomal locations), the presence of regions, either source or target, that interact with regulators led to an increase in the number of these that perform local search in that position. That the TF density around the target is so high in Figure 3B, in relation to the source, is due to close proximity effects. Indeed, the target was simulated to be 500bp away from where TFs are expressed, which is within the local (source) search (Figure 3A). The further away the target is moved, the less this second-timescale local search would be appreciated.

### Space as a design parameter

In previous results (*33*) we experimentally measured the effects of intergenic distance in a positive regulation (i.e. induction) using components of the TOL pathway (*49*) of the environmental bacterium *Pseudomonas putida* (*50*, *51*). Specifically, the gene *xylS* (source), which expresses XylS regulators was inserted in proximity to, or separated from, the promoter *Pm* (target), which is in turn activated by XylS. Spatial effects were quantified by measuring expression of *Pm-gfp* fusions in single cells with flow cytometry (Figure 4A). Results suggested that space could be used as a design parameter for selecting output levels, since the performance of the regulatory circuit changed according to spatial configuration. Specifically, when the source-target distance was minimized, reporter expression was fully *on* (i.e. narrow distribution to the right of the plot). In contrast, when such distance was increased, the expression became very noisy (i.e. wide distribution from left to right). Here, we compared the expression distributions in the experimental data with simulations by modulating source-target distance in our spatial model. As it can be observed in the side-by-side comparison of Figure 4A, the model for spatial regulation presented here gave an accurate reproduction of the experimental information - something that was not possible with homogeneous (i.e. non-spatial) models.

**Figure 4:**
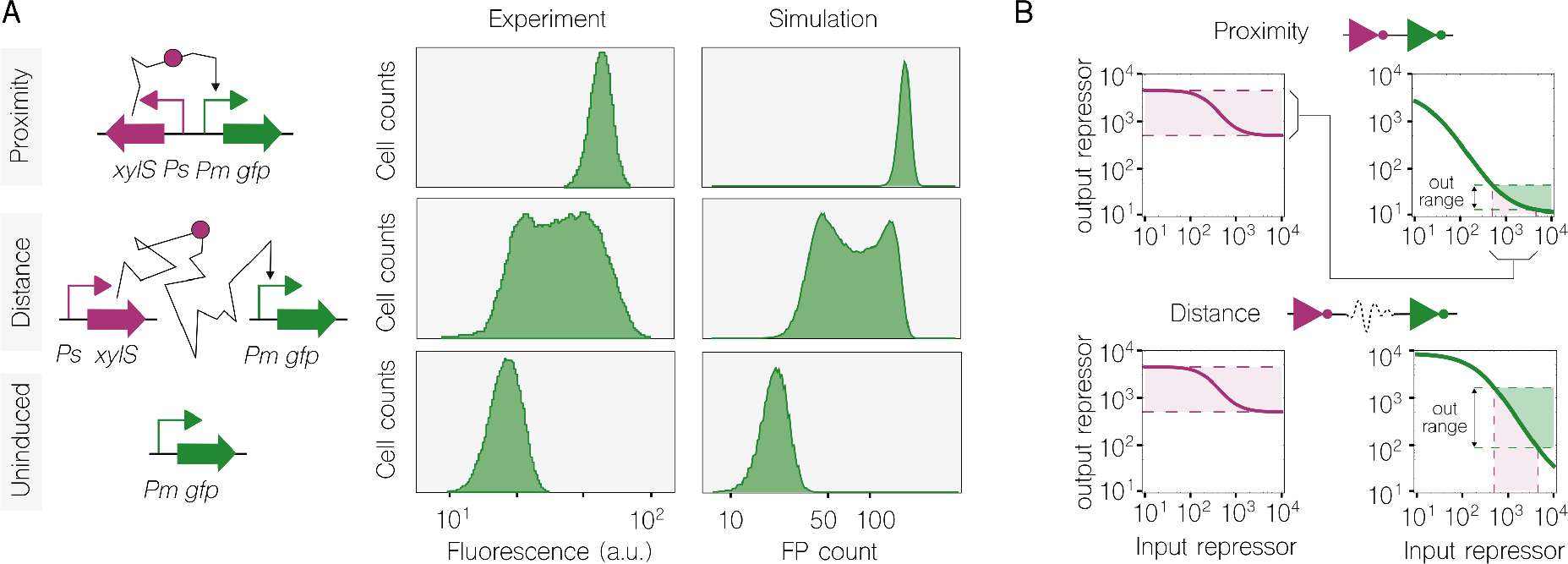
Space as a design parameter. **(A)** Reproducing the experimental results from Goñi-Moreno et al.(*33*) with the proposed spatial model. These experimental results measured gene expression noise in a construct where the source (gene *xylS*) of transcription factors (XylS) and its cognate target promoter (*Pm*) were engineered to be in either [i] proximity or [ii] in distant locations inside the bacteria *Pseudomonas putida*. In vivo flow cytometry results (adapted from (*33*)) and their in-silico counterpart look alike by modulating the intergenic distance alone. **(B)** Two genetic inverters modify their input-output mapping when either co-located or spatially separated. Top graphs show the simulation of two genetic inverters connected (i.e. the output of the first regulates the input of the second) in proximity. The output of the second inverter shows a very limited, and rather low, dynamic range (green shaded region). Bottom graphs show the same simulation but changing the location of the inverters to be far apart. The connection changes and the dynamic range at the ouput increases. Both examples suggest that space can be effectively used as a design parameter.

Moreover, the model shed light over mechanistic details that were difficult to elucidate on the experimental setup. For instance, the experimental construct did not offer quantitative separation metrics in the case of the distant scenario, since the source was inserted in a single-copy (mega)plasmid and the target in the chromosome. Being that the plasmid was located at the centre of the cell (*33*) it is safe to assume a middle distance following the scale shown in Figure 2E. Upon fitting the experimental data with the spatial model, separation was found out to be around 2kbp. In the case of co-localisation, both source and target were inserted in the same chromosomal location (*attTn7*), thus separation should not exceed tens of base pairs. This distance was set up to 100bp in the model.

Finally, we illustrate the potential of space as a design parameter in the context of genetic combinatorial circuits. In these, information is transmitted in the form of digital-like values, 0/low expression and 1/high expression, through regulatory interactions. Each of the components of a circuit can be seen as an electronic device that gets a value in the input and returns a value in the output after some processing. In order to get optimal performance, a key feature for a circuit is that the output of a component must be *compatible* with the input of the next one. Compatible in that the first component’s output range (i.e. the gap between the lower and higher values that can be produced as output) must be sufficient to differentiate two distinct values at the input of the second component. Otherwise, that connection will not be able to propagate digital values.

Currently, the lack of such compatibility between components is solved by simply selecting different components. That is, a given genetic logic gate can be replaced by another one if it is not ideal for a specific regulatory interaction. This approach is followed by, for instance, the tool Cello (*7*), a design automation platform that allows a user to turn high-level specifications into the DNA sequence of the corresponding genetic circuit. However, this procedure is limited by the catalogue of available components that could be used to this end - which is, in most of the cases, just a handful. Our take here is that there would be no need to replace the components if optimal performance could be achieved by organizing them into a different configuration within the volume of the cell. Figure 4B shows the compatibility between two theoretical genetic inverters, which are components that *invert* the input signal: if the input is 1, the output would be 0, and vice-versa. As can be observed, by solely changing the inverters from co-located to spatially separated, the output range of the second one improved substantially. Importantly, it was only distance that was modified in the model; the rest of the parameters in the simulation were the same. This suggests that the catalogue of *functions* that could be used to replace circuit components may not only be formed by different DNA devices, but by the same ones with different spatial arrangement.

## 3 Discussion

The rational design of gene regulatory circuits builds on our mechanistic understanding of gene regulation. Such understanding is commonly captured by mathematical models, that allow for turning mechanistic details into design principles. Although well-studied, gene regulation is still based on unclear dynamics; at least, not clear enough for the rigorous mathematical formalisation that modelling needs. Here, we focus on the implications that the separation between genetic components, within the chromosome, has on their final performance.

Ever since advances in technology allowed for it, the interest on the intracellular spatial organisation of the regulatory machinery has increased (*25* –*28*, *52*). However, current descriptions of how regulatory interactions are affected by intergenic distance are still somewhat controversial. It is the case that similar experimental setups had shown both significant (*29*) and insignificant (*53*) effects of such chromosomal separation. Our previous experimental work (*33*) showed not only a decrease in gene expression, upon inserting the regulator source *far* away, but also an increase of the noise pattern. Therefore, it is both expression and noise that were suggested to be modified by solely altering the spatial configuration of the regulation. There could be many potential causes of these conflicting results, such as growth rates, bacterial species/strains, genetic machinery and measuring methods. Therefore, the formalisation of such dynamics in a modelling framework is essential; model-guided analysis of *spatio-temporal* gene regulation may turn decisive to elucidated these details. Although existing tools such as SmolDyn (*54*), eGFRD (*55*), SMeagol (*56*) and others (*57*) simulate spatial dynamics to various extents, they do not focus on gene regulatory circuits, which is the goal of our study.

Our model builds on a previous definition of *facilitated diffusion* (*36*) that include the classification of transcription factors (TF) into two groups: of *local* and *global*. This is a crucial feature of the model. While local TFs perform a one-dimensional search along the chromosome close to their coding gene, global TFs undergo rapid three-dimensional diffusion through the cytoplasm. We adapted this definition to include the specific dynamics of gene expression (i.e. transcription and translation) and gene regulation (i.e. TF-promoter interplay). By considering the distribution of TFs along the chromosome (due to diffusion) and the binding rates of the system, we calculated the time it takes for a TF to bind its cognate promoter at any given location. While this model was able to accurately reproduce experimental observations (see Figure 4A) (*33*) (which seems not to be possible with *non-spatial models*), we identified two areas of improvement that will be the focus of successive studies. Firstly, the model fails to simulate Hill function-based dynamics. This is a direct consequence of diffusion, since it makes it very challenging to calculate cooperativity. Secondly, the model does not account for the three-dimensional folding of the chromosome. Rather, it assumes a homogeneous shape along the cell. However, since Hill functions are a phenomenological description, rather than a physical model, and nucleoid structure is not rigorously known, none of these features would make the model physiologically more accurate.

The growing availability of methods for chromosomal insertions (*58* –*60*), allows researchers to transit from plasmid-based to chromosomal-based synthetic systems. While plasmids are metabolically demanding for bacteria (*61*), chromosomal insertions have been found to be more efficient (*62*). As increasingly precise responses are required from synthetic constructs, each component may need a specific location within the chromosome for optimal performance - an issue that deserves further attention. We advocate for the use of inter-component distance as a design parameter for synthetic circuits, and the use of spatiotemporal modelling in order to establish the engineering principles of such three-dimensional design.

## 4 Methods

### A facilitated diffusion model of gene regulation

We develop on previous models of *facilitated diffusion* dynamics (*36*, *63*) that crucially add the space-dependent 1D local search (*64*) to the space-independent 3D diffusion of molecules (*64*). The present model adds, on top of this the mechanistic dynamics of gene expression and gene regulation. While the former uses single rates e.g. transcription and translation, the latter involves two particles (e.g. a TF binds a promoter) and are described by the combination of a rate and a concentration. Every reaction that involves two particles (see assumptions below) was calculated by explicitly modelling diffusion.

### Assumptions taken by the model

A number of simplifications were considered: [i] transcription and translation are single-particle reactions i.e. polymerases and ribosomes are not explicitly modelled; [ii] transcription factors perform a local search upon translation; [iii] transcription factors only bind to DNA during 1D search; [iv] global search is a combination of 3D and 1D diffusion i.e. TFs slide many times during global search (*39*, *65*, *66*); [v] the chromosome is not explicitly modelled (DNA folding not considered) which reduces the problem from solving a 3D reaction to solving a 1-dimensional one;[vi] transcription and translation are co-localized, as it is generally accepted for prokaryotes (*27*); [vii] TF polymers are formed directly after translation and there is a high chance they bind to the chromosome at that moment (these mechanistic details are still unclear).

### Calculation of TF distribution along the chromosome

The distribution of TFs that corresponds to global search is uniform i.e. the density of global TFs is constant throughout the chromosome. This is due to the rapid 3D diffusion of TFs within the volume of the cell; once a TF leaves the initial local search, it can be anywhere. Specific unbinding rate (i.e from the target into the cytoplasm) was set to 1/200 *s*^−1^ (*67*), which corresponds to an experimental *absolute* value that already accounts for different specific TF dynamics, like re-binding (*68*) (i.e. fast binding after unbinding), for which metrics are not well established. The uniformity of the global search draws on such unbinding rate.

Local search, however, is explicitly modelled. Its corresponding local distribution depends on the 1D diffusion constant and the rate of non-specific unbinding (i.e from a non-target DNA location into the cytoplasm). In order to arrive at such distribution, we firstly solve the following equation that describe the density of TFs in time and space:

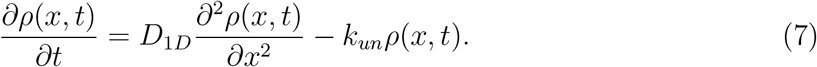

where *t* = 0 is the time of TF expression, *x* = 0 is the location of the source gene, *ρ* is the TF density, *D*_1*D*_ is the 1D diffusion rate and *k*_*un*_ is the unspecific DNA unbinding rate. Equation 7 is three-dimensional (*ρ*, *t*, *x*); the integration of *t* from 0 to *∞* returns the distributions of Figure 3B:

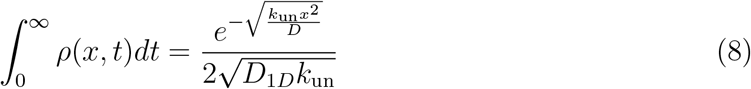

### Calculation of search times

The local and global search times define the average time it takes to a TF to find the target promoter. In the case of global TFs, given that their distribution along the chromosome is uniform, this value is calculated by:

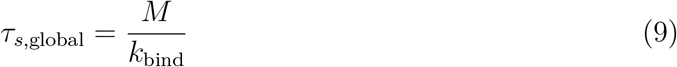

where M denotes the length of the chromosome. Distance from the source does not play a role in the calculation of global search since, as stated above, 3D diffusion is so fast that spatial autocorrelation with the source is quickly lost. Local search time, however, does consider the separation (in bp) from source. Along with the binding rate, the distribution that results from Equation 8 generate the local search times. The derivation (Supplementary File S1) returns the following:

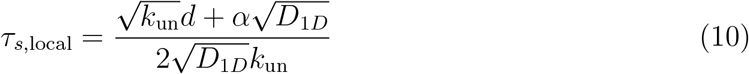

where the parameter *α* (0 < *α* < 1) depends on *D*_1*D*_ and *k*_un_ (Supplementary File S1) and describe the effect of re-binding (binding immediately after unbinding). At *α*=0 the TF would never leave the target once it finds it.

### Binding rates

The homogeneous model of Figure 2 uses a binding rate of 10^6^ *M*^−1^*s*^−1^. Spatial-based binding rates, however, do not follow the same metric scale. In the spatial model we separated the binding rate into two different values: the time it takes for a TF to find the target (i.e. local/global search times), which depends on diffusion and unbinding rates, and the propensity (*γ*, dimensionless) of a TF to bind after it finds the target. Figure 2D shows the effect of decreasing this parameter tenfold. Unbinding rates are not affected by spatial constraints, thus obtained from literature (see Table 1).

### Stochastic Simulations

For the stochastic simulations in Figures 2B-E and 4A reactions were implemented in the Gillespie algorithm (*72*).

### Homogeneous model

To compare with our model we use a homogeneous (i.e. non-spatial) model. This model uses the same rates for transcription, translation and degradation than the spatial one, since these are not affected by distance. The differences are that [i] the TFs are not classified into local and global, and [ii] binding rate is measured in *M*^−1^*s*^−1^ (Table 1).

## Acknowledgement

This work was supported by the SynBio3D project of the UK Engineering and Physical Sciences Research Council (EP/R019002/1) and the BioRoboost Contract of the European Union (820699).

