## Supplementary File S1 for "A model for the spatio-temporal design of gene regulatory circuits"

### Supplementary Information

Ruud Stoof, Alexander Wood, and Ángel Goñi-Moreno*

*School of Computing, Newcastle University, Newcastle upon Tyne, NE4 5TG, UK.*

#### 1 Quasi steady state mRNA

The system of Ordinary Differential Equations was simplified using the quasi-steady-state assumption for mRNA:

$$m_i^*(P_i) = \frac{\beta_{\text{transc}}(P_i + \alpha_{\text{leak,i}})}{k_{\text{Dm}} + \mu} \tag{S1}$$

replacing equations 3 and 6 with:

$$\frac{dR_{\text{local}}}{dt} = \beta_{\text{transl}} m_R^*(P_R) - (k_{\text{Dp}} + \mu)R_{\text{local}} - k_{\text{unb}} R_{\text{local}} - \frac{P_{\text{gfp}} R_{\text{local}}}{\tau_{\text{s,local}}} \tag{S2}$$

$$\frac{d\text{FP}}{dt} = \beta_{\text{transl}} m_{\text{gfp}}^*(P_{\text{gfp}}) - (k_{\text{Dp}} + \mu)\text{FP} \tag{S3}$$

#### 2 Derivation of $\tau_{\text{local}}$

To get the local search time at a specific inter-genic distance we solve Equation7:

$$\rho(x,t) = \frac{e^{-\frac{x^2}{4Dt} - k_{\mathrm{un}} t}}{2\sqrt{\pi}\sqrt{Dt}} \tag{S4}$$

Now we calculate the average time it takes for the TFs to bind within this limit:

$$\frac{\int_0^\infty t k_{\mathrm{bind}} \rho(x,t) dt}{\int_0^\infty k_{\mathrm{bind}} \rho(x,t) dt} = \frac{\int_0^\infty t \rho(x,t) dt}{\int_0^\infty \rho(x,t) dt} = \frac{\sqrt{k_{\mathrm{un}}}\, |x| + \sqrt{D}}{2\sqrt{D} k_{\mathrm{un}}} \tag{S5}$$

If this is compared against the solution $\frac{|x|}{2\sqrt{Dk_{\mathrm{un}}}}$ in Wunderlich and Mirny (*1*), we see that there is an extra term of $\frac{\sqrt{D/k_{\mathrm{un}}}}{2\sqrt{Dk_{\mathrm{un}}}}$, referred to as $\alpha$. This parameter defines the effect of non-immediately binding. Let $0 < \alpha < 1$ , $\lim_{k_{\mathrm{bind}} \to \infty} \alpha = 0$ and $\lim_{k_{\mathrm{bind}} \to 0} \alpha = 1$.

By substituting we obtain:

$$\tau_{\mathrm{local}} = \frac{d + \alpha\sqrt{D/k_{\mathrm{un}}}}{2\sqrt{Dk_{\mathrm{un}}}} \tag{S6}$$

where we fill in the distance between the source and target, $d$, for $|x|$.

Solving the binding rate from the global search time returns a value for a TF and a location to be ca. $5000 s^{-1}$ (using the parameters $M$ and $\tau_{\mathrm{global}}$), hence $\alpha \approx 0$.
